# Context-dependent coordination of movement in *Tribolium castaneum* larvae

**DOI:** 10.1101/2024.06.17.598650

**Authors:** Bella Xu Ying, Maarten F. Zwart, Stefan R. Pulver

**Affiliations:** Institute of Behavioural and Neural Sciences, University of St Andrews, St Andrews, United Kingdom; Centre of Biophotonics, University of St Andrews, St Andrews, United Kingdom; School of Psychology and Neuroscience, University of St Andrews, St Andrews, United Kingdom

**Keywords:** *Tribolium castaneum*, locomotion, kinematics, motor control, neuroethology, central nervous system

## Abstract

Insect pests, like the red flour beetle *Tribolium castaneum*, destroy up to 20% of stored grain products worldwide, making them a significant threat to food security. Their success hinges upon adapting their movements to unpredictable, heterogeneous environments like flour. *Tribolium* is well developed as a genetic model system; however, little is known about their natural locomotion and how their nervous systems coordinate adaptive movement. Here, we employed videographic whole-animal and leg tracking to assess how *Tribolium* larvae locomote over different substrates and analyze their gait kinematics across speeds. Unlike many hexapods, larvae employed a bilaterally symmetric, posterior-to-anterior wave gait during fast locomotion. At slower speeds, coordination within thoracic segments was disrupted, although intersegmental coordination remained intact. Moreover, larvae used terminal abdominal structures (pygopods) to support challenging movements, such as climbing overhangs. Pygopod placement coincided with leg swing initiation, suggesting a stabilizing role as adaptive anchoring devices. Surgically lesioning the connective between thoracic and abdominal ganglia impaired pygopod engagement and led to escalating impairments in flat-terrain locomotion, climbing and tunnelling. These results suggest that effective movement in *Tribolium* larvae requires thoracic-abdominal coordination, and that larval gait and limb recruitment is context-dependent. Our work provides the first kinematic analysis of *Tribolium* larval locomotion and gives insights into its neural control, creating a foundation for future motor control research in a genetically tractable beetle that jeopardizes global food security.

**Summary statement:** Red flour beetle larvae walk with a legged wave gait and use their tails as anchors to climb inclines and tunnel into flour.

## Introduction

Stored product insects are devastating agricultural pests. These animals have been estimated to cause 9% of postharvest loss in developed countries and 20% in developing countries (reviewed in (Phillips and Throne, 2010)). The red flour beetle, *Tribolium castaneum (Tribolium),* is representative of many stored product pest insects presenting a major challenge to global food security. Larvae and adults consume wheat products like flour by efficiently moving over and tunnelling through these unstable, shifting substrates (Good, 1936). Therefore, understanding how *Tribolium* adapt their movements to heterogeneous and challenging terrain could provide insights into conserved principles of motor control whilst aiding the development of new pest control approaches.

Animals use various strategies for adapting movement to the environments in which they live, continuously monitoring and adjusting movements on a cycle-by-cycle basis to cope with unexpected obstacles and/or drastic variations in the physical properties of 3-dimensional terrain. Studying the neural control of movement adaptability has been a long-standing focus of motor systems research across multiple species. Terrestrial animals with internal, external, or hydrostatic skeletons have all evolved different strategies for movement control in different contexts (reviewed in insects (Mantziaris et al., 2020; van Griethuijsen and Trimmer, 2014), mammals (Leiras et al., 2022) and birds (Daley, 2018)). Some animals deploy limbs (Hooper et al., 2009; Picker et al., 2012; Triphan et al., 2010), some utilize soft outer cuticular features (Digumarti et al., 2019; Booth et al., 2024) and others adapt body parts not typically involved in locomotion to solve motor problems (Shield et al., 2021). Similarly, *Tribolium* exhibit a diverse behavioral repertoire in response to different sensorimotor cues, and adult anatomy has been well-described. Previous work has shown that *Tribolium* can move through a variety of stored grain, but prefer finely grained, whole wheat flour over coarser bran (Đukić et al., 2020). Moreover, adults can navigate in response to attractive and aversive odors (Đukić et al., 2021; Gerken et al., 2018), shapes (Semeao et al., 2011), light levels (Duehl et al., 2011), colors (Prakashini et al., 2023), and different habitat cues (Romero et al., 2010). Both larvae and adults have articulated, jointed limbs (Good, 1936), with larvae being composed of both soft ventral cuticle and semi-hardened dorsal cuticle. While anatomical studies have begun to characterize the larval neuromuscular system (Schultheis et al., 2019), knowledge concerning the specific kinematics of locomotion and how the *Tribolium* nervous system recruits thoracic limbs and abdominal segments to meet different locomotor demands is extremely limited.

Past studies have prioritized adult anatomy and behavior, although larval instars are equally capable of maneuvering and colonizing stored grain. Here we characterize locomotor parameters in late-instar *Tribolium* larvae to gain insights into how these animals adapt movements to their environment. We analyze how larvae negotiate different substrates and combine whole-animal and leg tracking to characterize and quantify leg kinematics underlying a range of locomotor speeds. We find larvae have an ethologically relevant preference for fibrous substrates and walk primarily with a posterior-to-anterior-propagating bilaterally symmetric gait, which shifts towards uncoordinated exploratory movements at slower speeds. Severing the connectives between thoracic and abdominal segments impairs walking, climbing, and tunneling but does not disrupt fundamental leg kinematics. In addition, we characterize a pygopod planting behavior, abolished in animals with severed connectives, which is used for stability and propulsion under different motor contexts (challenged locomotion vs tunnelling). These results provide first insights into the biomechanical underpinnings of *Tribolium* movement and the neural basis of ethologically relevant motor programs in an insect species with direct impacts on global food security.

## Materials and methods

### Animals

Wild-type *Tribolium castaneum* (San Bernardino strain) were reared in 500ml borosilicate bottles with perforated lids (VWR SCOT291182809) at 29°C in organic whole wheat flour and 5% dried active yeast per weight. 4-5^th^ instar larvae approximating 4-6mm in length were used in all experiments.

### Statistics

Main text results and box plots are given as mean ±SE, and box whiskers are ±SD unless otherwise stated. Details of statistical analysis are included under each experiment’s subsection below and in figure legends.

### Substrate testing

Basic locomotor ability was tested on twelve substrates – white printer paper, cardboard, parafilm, lens paper, Sylgard™, glass, velvet, sandpaper, plastic, packing foam, white flour, whole wheat flour, 1%, 5% and 10% agarose. Ten larvae were placed in the center of the substrate and recorded for 1 minute. Movies were recorded with a mobile camera (SM-A415F/DSN, Samsung, Seoul, South Korea) at 30fps and analyzed with Fiji (Fiji is Just ImageJ) (Schindelin et al., 2012). Larval head position was manually tracked for the first 15 seconds (since larvae on preferred substrates often walked out of frame within this time window) with the Manual Tracking plugin (https://imagej.net/ij/plugins/track/track.html) for Fiji.

### Free locomotion assay

30 larvae were placed in groups of 3-4 in a 18x18cm arena lined with white printer paper and recorded for 2 minutes in darkness with a custom-built imaging setup using an MQ013MG-E2 XIMEA CMOS (Ximea GmbH, Münster, Germany) camera with a 35mm C-mount lens (Thorlabs Inc, NJ, USA). Animals were illuminated with 850nm infrared light using a Thorlabs M850L3 LED (Thorlabs Inc, NJ, USA). Larvae were tracked with FIMTrack (Risse et al., 2014). Statistical analysis on biases in headsweep direction was performed as an intercept-only general linear model (GLM) in R.

### Gait characterization

Ten larvae were placed on white printer paper under an Olympus SZX10 (Olympus Microscopes, Tokyo, Japan) dissection microscope at 0.8x magnification using a 0.5x objective. Animals were recorded with an MQ042RG-CM XIMEA CMOS (Ximea GmbH, Münster, Germany) camera at 60fps. 3-5 straight walk cycles were extracted and manually tracked with Fiji’s Manual Tracking plugin. Gait dynamics and locomotor parameters were analyzed with Python scripts for individual legs; cycle period was calculated as 1 over average cycle frequency. A Savitzky-Golay low-pass filter was applied to manually tracked velocities to discretize swing durations. Stance duration was calculated as the period between consecutive swings. Duty cycle was calculated as average swing/stance duration per segment over average cycle period per animal. Intrasegmental phase difference represents the phase difference between left and right legs within each thoracic segment. Intersegmental phase difference represents phase difference of left legs in each segment relative to left T3.

Coherence and phase analysis was performed with MATLAB application *Groundswell* (https://github.com/JaneliaSciComp/Groundswell). A fast Fourier transform was performed on the anchor leg’s velocity trace to determine its spectral composition. The anchor’s fundamental frequency was determined as the frequency with greatest power density (i.e., during swing phases) in the power spectrum. If the highest power frequency was ambiguous (i.e., appeared as blocks of plateaus on the spectrum), the middle value within the plateau was chosen, and the higher frequency block was used in cases with two adjacent blocks. Coherence and phase relationships of the other five legs to the anchor was compared at the determined fundamental frequency. *Groundswell* uses coherence magnitude as a measure of the linear relationship between two waveforms at the determined fundamental frequency, and calculates a coherence magnitude threshold with the formula:

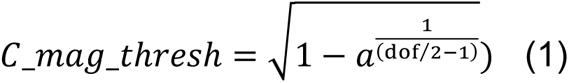

Where degrees of freedom (dof) = 2*R*K, in which R is the number of observations of the waveforms together and K is the number of tapers. This threshold determines the minimum coherence magnitude required for significant coherence between two signals with α = 0.05.

### Preliminary incline assay

Ten larvae were individually placed on custom-built platforms of varying incline (180° – flat substrate, 135° – slight upwards incline, 90° – vertical wall, 45° – overhang). Locomotion was filmed with an MQ022RG-CM XIMEA CMOS (Ximea GmbH, Münster, Germany) camera for 1 minute at 30fps with a Thorlabs OSL1 Fiber Illuminator (Thorlabs Inc, NJ, USA) with gooseneck lights at lowest intensity. Raw videos were optimized for FIMTrack in Fiji with custom-written macros. Statistical analysis on mean instantaneous velocity was performed as one-way ANOVA. Path curviness was calculated with the formula:

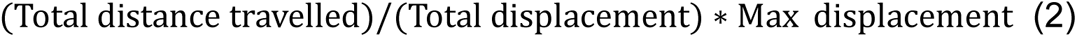

### Connective severing assay

In preliminary control assays, six larvae were anesthetized on ice and sequestered between 0.1mm tungsten pins ventral side up on a dissection dish. A cut was made with fine microdissection scissors on the cuticle line separating A1 and A2 segments. The blades were inserted at a ∼45° angle to the cuticle and pulled upwards to sever the connective. Sham larvae were given lateral lesions on A1. Control larvae were not lesioned but anesthetized on ice for equal time. Surgeries were verified via fillet microdissection post-experimentally. Larvae without clear, visible connective severing were discarded from results. Behavior was filmed with the setup outlined in *<u>Free locomotion assay</u>* 2 hours after surgery; raw videos were processed in Fiji and tracked with FIMTrack. Statistical analysis on mean instantaneous velocity was performed as a one-way ANOVA.

For following incline and tunnelling experiments, surgeries were repeated on seven larvae; sham animals’ lateral lesions alternated between left and right sides to minimize biases in turning. Different animals were used for incline and tunnelling assays to avoid excessive fatigue on lesioned animals.

### Top-down connective severing incline assay

Same methodology as *<u>Preliminary incline assay</u>,* excluding 90° due to short lifespans of severed animals. 2 larvae (1 control and 1 severed) could not be tracked with FIMTrack, thus mean velocity was obtained with wrMTrck (https://www.phage.dk/plugins/wrmtrck.html) and their tracks are therefore absent in Fig. 5A. Statistical analysis on mean instantaneous velocity was performed as one-way ANOVAs within each inclination with Bonferroni corrected post-hocs.

### Up-close connective severing incline assay

Control, sham and severed larvae were individually placed onto a walled 70mm x 3mm paper platform on a custom-built rig; incline was adjusted with a micromanipulator and verified with a protractor. Locomotion was filmed until the animal walked out of view with a MQ042RG-CM XIMEA CMOS (Ximea GmbH, Münster, Germany) camera connected with a 1”-32 C-mount (Thorlabs Inc, NJ, USA) to a Canon FR1048KWN lens (Canon Inc, Tokyo, Japan). If the animal remained immobile, filming was stopped at the experimenter’s discretion. 180° videos were filmed at 100fps and illuminated by an overhead Thorlabs M850L3 850nm light (Thorlabs Inc, NJ, USA). 45° overhang trials were illuminated from the bottom with a Stemmer Imaging CCS LDL100 850nm light (Stemmer Imaging Ltd, Tongham, UK) and limited to 50fps by the field of view. 135° and 90° were not tested due to the short lifespans of severed animals. For analysis, velocity was tracked with FIMTrack; pygopod planting events and start and stop frames of leg swings were manually tracked with Fiji’s Manual Tracking plugin. The constricted arena walls promoted exploration, therefore to effectively analyze locomotor parameters and gait kinematics, walk cycles following the T3 to T1 sequence were extracted and stance durations above median ± 1SD were discarded to filter out pause periods. Mean percentage of sequences extracted out of total tracked time (including pauses) per animal were: 55.4% (control 180°), 47.2% (sham 180°), 34.2% (severed 180°), 28.3% (control 45°), 24.9% (sham 45°) and 26.0% (severed 45°). Cycle period, stance duration and duty cycle retained their definitions as in *<u>Gait characterization,</u>* but swing duration was calculated as the time between swing start and swing stop frames.

A pygopod plant event was defined as a visible extension of the pygopods, forming almost a blunt triangular point, into the substrate, regardless of whether the abdomen is in contact with the substrate. The Freedman-Diaconis rule was used in Fig. 5R, using the formula:

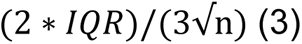

to calculate x-axis bin length. Planting rate was used for Fig. 5T to normalize for variable video lengths, calculated as number of events over total tracked time. Pearson’s correlations were applied for regression analyses.

### Tunneling assay

Post-surgery animals were allowed to wake and begin walking on paper before testing. A circular dish was filled with ∼10mm of flour; control, sham and severed larvae were individually placed and filmed with an MQ0113RG-E2 XIMEA CMOS (Ximea GmbH, Münster, Germany) at 15fps under a Nikon SMZ745T dissection scope at 0.63x magnification. Filming stopped at 2 minutes regardless of how deep larvae tunneled. Body lengths at time 0s and 120s were measured in Fiji. Statistics was performed as a Kruskal-Wallis test and Dunn post-hoc analysis with Bonferroni correction (given non-normal data distribution).

## Results

### *Tribolium* larvae locomote most efficiently over fibrous substrates

*Tribolium* beetles take approximately 35 days at 29°C to undergo complete metamorphosis, comprising four distinct developmental stages – egg, larval, pupal, and adult (Devi and Devi, 2015) (Fig. 1A). The larval stage is characterized by three thoracic segments, each supported by a pair of legs, and nine abdominal segments. Larvae possess two small, rounded ventral pygopods in the most posterior abdominal segment, and two spiked urogomphi dorsally (Fig. 1B).

**Figure 1:**
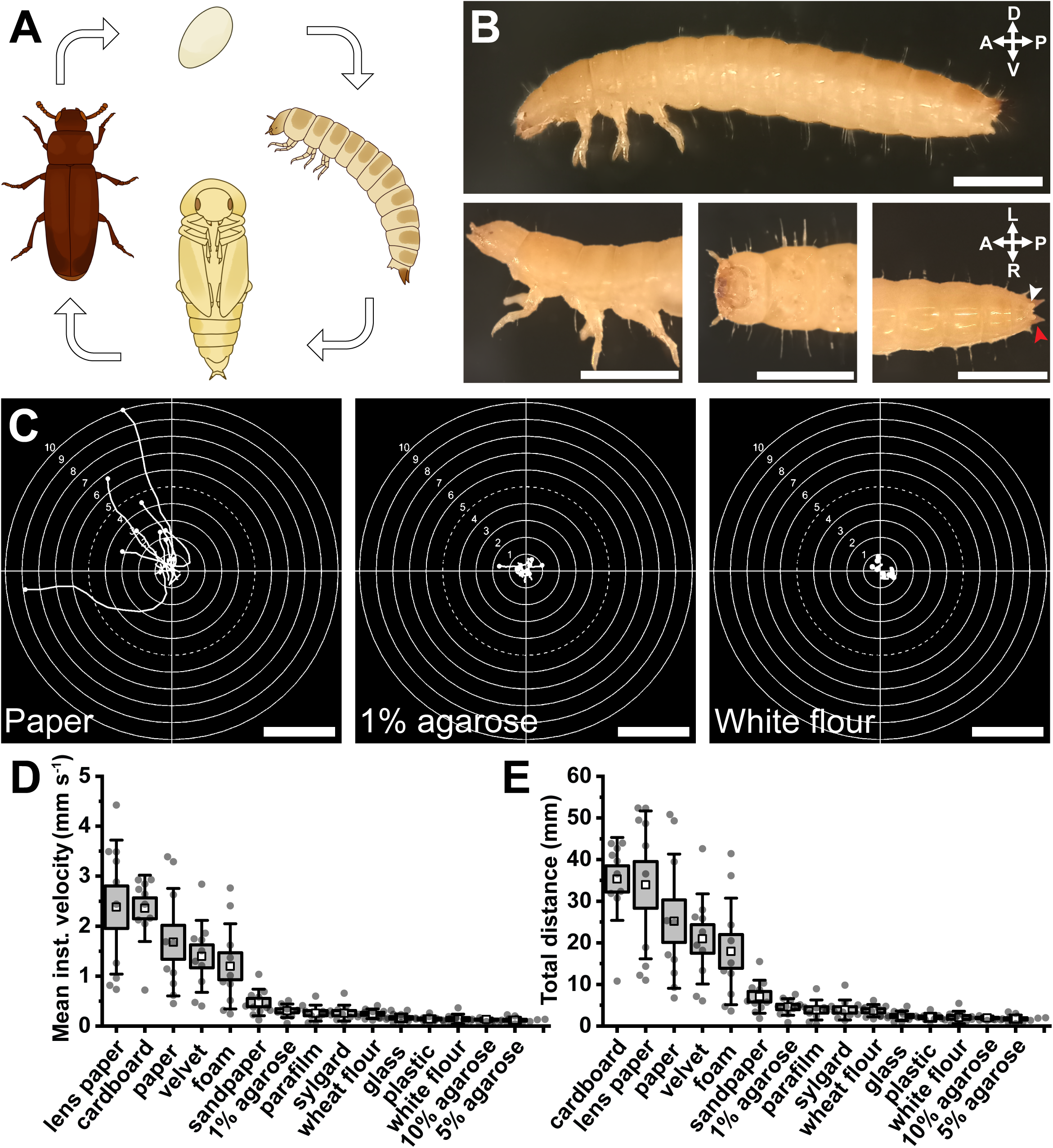
*Tribolium castaneum* larval locomotion is most efficient on fibrous substrates. (A) Diagrammatic representation of *Tribolium castaneum* life cycle. (B) Morphology of 4^th^-5^th^ instar larva under a 10x objective. Full body image at 2x magnification and insets at 5x magnification showing three thoracic leg pairs, nine abdominal segments, pygopods (white arrow) and urogomphi (red arrow). Scale bars = 1mm. (C) Larval paths over select substrates during first 15 seconds of free locomotion. Scale bar = 10mm. (D) Mean instantaneous velocity across fifteen substrates, ordered highest to lowest. (E) As (D), for total distance travelled. White squares = mean, boxes = mean ±SE, whiskers = ±SD overlayed on raw data. N = 10.

We first established larval substrate preference based on ease of locomotion. We found larvae crawl best on textured, fibrous, wood-based substrates such as paper, cardboard, and lens paper (Fig. 1C) <u>(Movie 1)</u>. In contrast, they had substantial difficulty gripping smooth surfaces like plastic and glass, thus preventing forwards momentum. Similarly, their legs slipped on or stuck to deformable substrates like Sylgard™ and agarose, also impeding coordinated progress (Fig. 1D,E). Notably, on these substrates, we observed pygopod planting for substrate engagement in an anchoring motion, which, when released, propelled the larvae forwards <u>(Movie 1)</u>. This abdominal engagement was rarely observed on fibrous substrates. Surprisingly, crawling was heavily impeded over whole wheat flour and white flour (Fig. 1C-E), as larvae attempted to tunnel upon contact with these substrates.

### Larvae exhibit diverse motor behaviors during free locomotion

Next, movement parameters during free locomotion were analyzed to quantify the spectrum of larval motor capabilities. Paper was used as the default substrate in subsequent experiments given larval preference, accessibility, and ease of imaging. *Tribolium* larvae exhibited heterogeneous walking patterns, and favored edge exploration, i.e., they rarely strayed from arena edges once they found them (Fig. 2A). Additionally, we observed persistent burrowing attempts at the paper edges during preliminary trials in which the paper substrate was not fully adhered to the arena platform.

**Figure 2:**
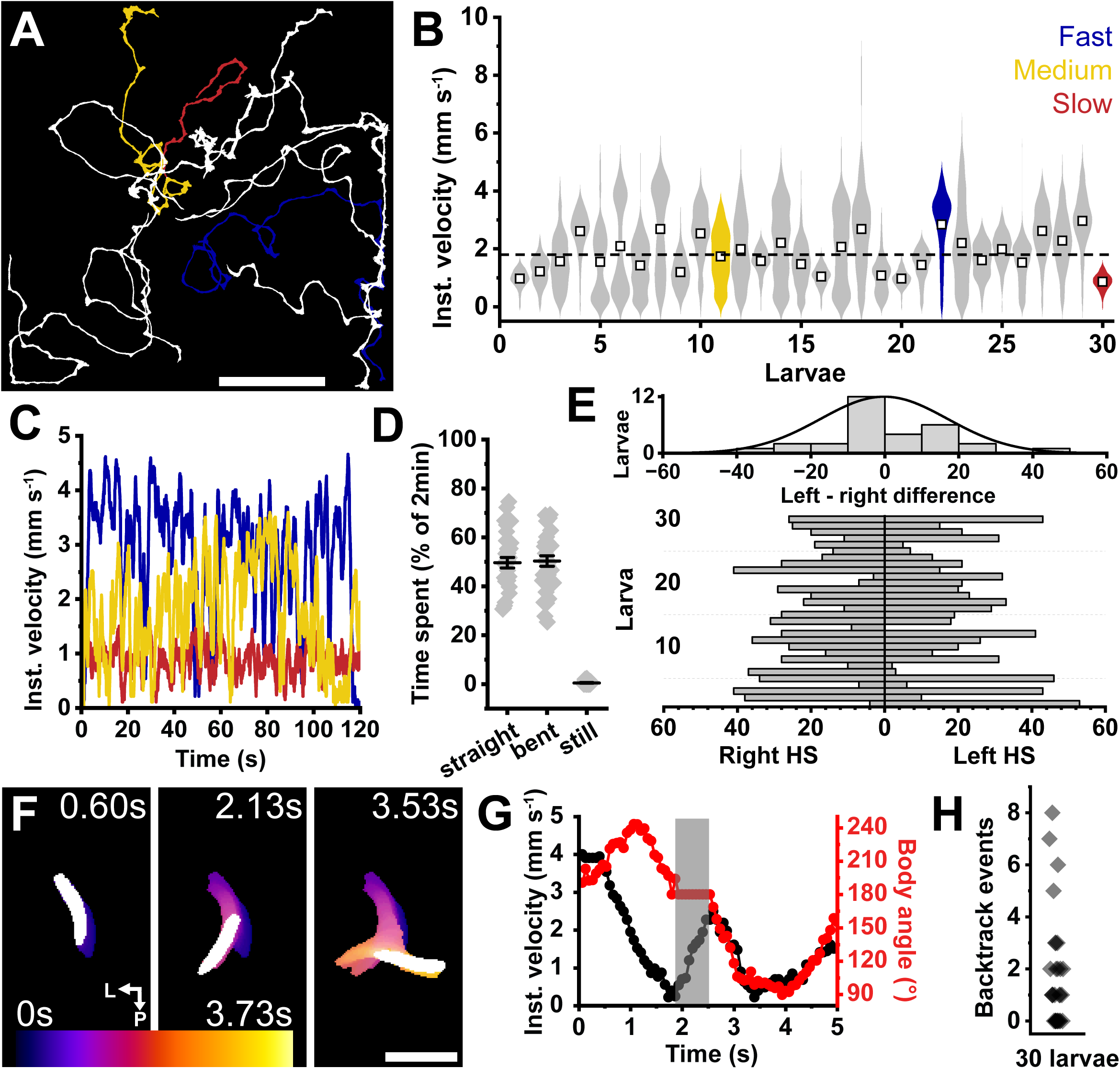
Larvae display inter- and intra-animal variability in free locomotion parameters. (A) Larval locomotor paths across white paper for 2 minutes. Colored paths represent fast (blue), intermediate (yellow) and slow (red) larvae. Example larvae were selected based on highest, lowest and intermediate mean instantaneous velocity out of 30 larvae. Scale bar = 5cm. (B) Violin plot showing intra-animal variability in instantaneous velocity. White squares = mean. (C) Instantaneous velocity traces across 2 minutes of colored animals in (A) highlighting inter- and intra-animal variability. (D) Proportion of time spent in distinct locomotor behaviors: straight = 180° body angle and >0mm s^-1^ velocity, bent = non-180° body angle and >0mm s^-1^ velocity, still = 0mm s^-1^ velocity regardless of body angle. Whiskers = mean ±SE, n = 30. (E) Bottom: total left and right headsweep events by individual larvae. Left headsweep = ≥210° body angle, right headsweep = ≤150° body angle as defined by FIMTrack. Top: histogram of headsweep directional difference (left minus right headsweep count); intercept-only GLM, *p =* .869 for hypothesis that left headsweep probability out of total is significantly different from 0. (F) 3 frame sequence during a backtrack and redirect behavior. Frames are color-coded for total time (3.73s); temporal larval trajectory is overlayed beneath current frame (white). Note the center frame showing more posterior body position relative to preceding frames, indicative of “backtracking”. Scale bar = 5mm. (G) Instantaneous velocity and bodybending angle against time during a single backtrack and redirect behavior sequence. Gray area highlighting backwards locomotion. (H) Total number of backtrack and redirect events during two minutes of free locomotion. N = 30.

Locomotion on paper had a mean instantaneous velocity of 1.8 ± 0.1 mm s^-1^ (range = 0.9 to 3.0 mm s^-1^; larval sizes ranged 4-6 mm), and animals displayed a spectrum of intra-animal velocity variability during only 2 minutes of exploration (Fig. 2B). Faster animals were not constantly fast, but instead varied speed throughout locomotion, whereas slower animals were limited to lower speed ranges (Fig. 2C). Larvae spent equal time walking straight (49.6%) and turning (50.4%) (Fig. 2D). Animals decelerated to turn but rarely stopped completely, thus staying immobile only 0.5% of the time (Fig. 2D). Finally, while some larvae showed a bias in turning direction, many engaged equally in left and right headsweeps during exploration (p = .869; Fig. 2E).

We observed a larval motor program we termed “backtrack and redirect” <u>(Movie 2)</u>, defined as a headsweep/turn in one direction, followed by visible backwards locomotion, and a contralateral headsweep/turn and forwards locomotion (Fig. 2F), which on average lasted 3-6 seconds. This behavior followed a distinct sequence of instantaneous velocity and body angle changes. First, velocity dropped as the animal turned (body angle deviating from 180°), followed by an increase in velocity at 180° body angle due to backtracking, and ended with another deceleration and deviation from 180° in the opposite direction from the animal’s original trajectory (Fig. 2G). While 63.3% of larvae displayed this motor program at least once during 2 minutes of free locomotion (Fig. 2H), some displayed it a maximum of 8 times, and a third never engaged in this behavior.

### Gait coordination and leg kinematics during locomotion is speed-dependent

To characterize natural gait, leg kinematics and the coordination between thoracic segments, we tracked legs during straight walking bouts on paper. We found *Tribolium* larvae walked with a bilaterally symmetric posterior-to-anterior wave gait (Fig. 3A). Then, we determined the frequency of locomotion for each animal by performing a Fourier transform on leg velocities and selecting the frequency with highest power of an anchor leg (left T1). We defined swing phases by a leg’s lifting and planting on the substrate and stance phases as the duration in which legs are in contact with the ground. An average cycle period was then calculated for each leg, then averaged per animal. After a swing, legs usually stanced next to the legs of the segment in front, and these often began their swing phase before the previous contacted the ground (Fig. 3A). This pattern was observed most prominently within the T2 to T1 transition as their swing phases overlapped more visibly (Fig. 3B-D).

**Figure 3:**
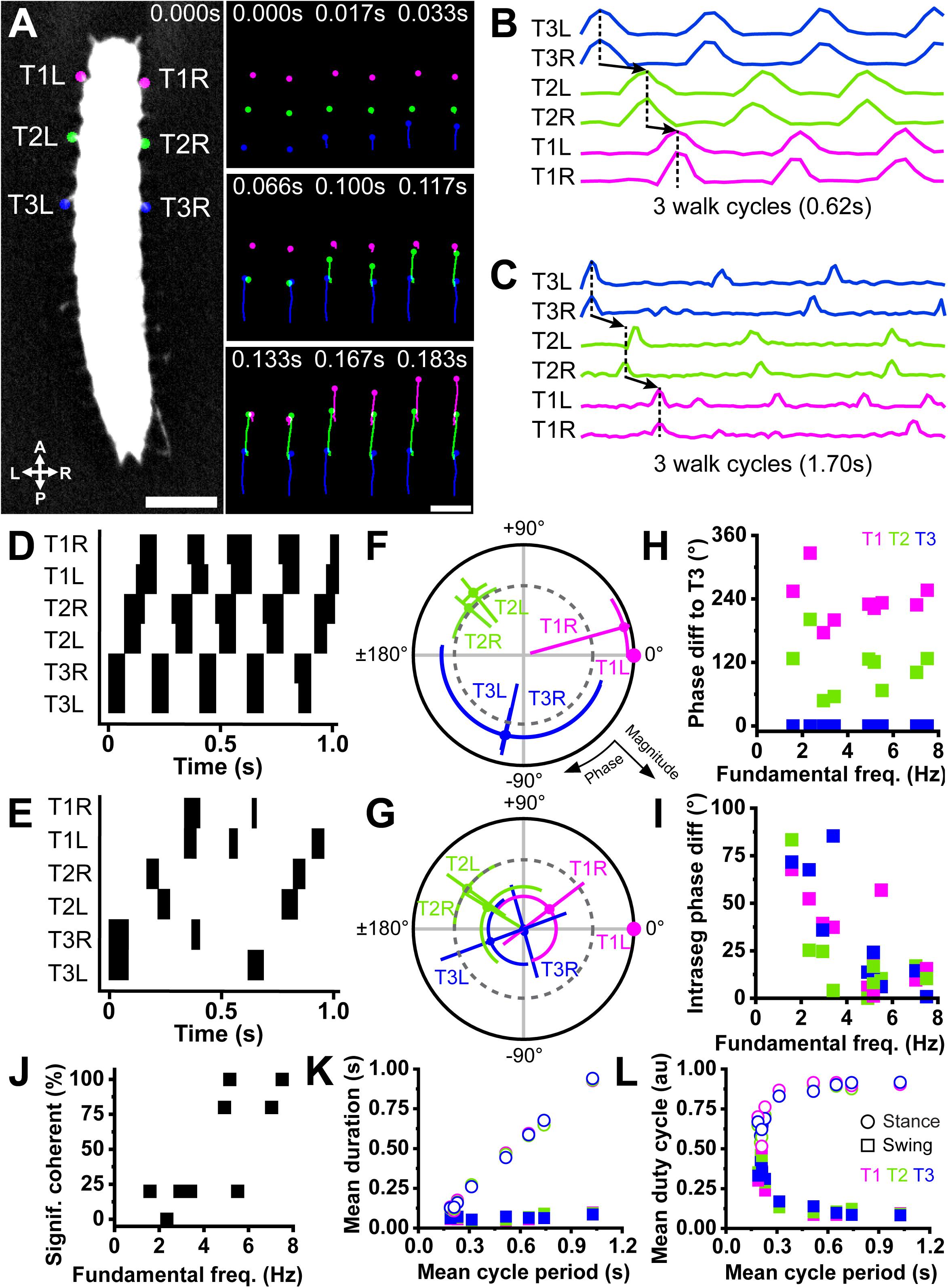
Gait coordination and leg kinematic coherence are largely speed dependent. (A) Left: example top-down view of *Tribolium* larva with labelled legs across three thoracic segments: T3 (blue), T2 (green) and T1 (magenta). Scale bar = 1mm. Right: select frames of tracked leg positions showing progression from T3 to T1 in one representative walk cycle. Note the bilateral symmetry as intrasegmental leg pairs swing. Scale bar = 1mm. (B,C) Example velocity traces of legs for (B) fast and (C) slow animals across three walk cycles. Note the greater intersegmental coordination in (B) than (C). (D,E) Swing (black) and stance (white) phases in the same (D) fast and (E) slow animals in (B,C). (F,G) Polar plots of coherence phase and magnitude of waveforms in (B,C) using T1L as anchor/0° for (F) fast and (G) slow animals. Negative phase values temporally precede positive ones. Dark grey dotted inner circle indicates α = 0.05 for coherence magnitude. (H) Scatter of phase difference between left legs of each segment and T3L across fundamental frequencies (proxy for walking speed). (I) Scatter of phase difference between legs of each segment. (J) Percentage of significantly coherent legs (out of 5) to anchor (T1L) per animal. (K) Mean swing and stance durations and (L) duty cycles for each segment (color-coded) across 3-5 walk cycles as a function of mean cycle period. N = 10.

Fast (Fig. 3B,D,F) and slow (Fig. 3C,E,G) walk cycles differed markedly. In faster animals, typical walk cycles averaged 0.2 ± 0.007s and were initiated by a T3 swing phase, followed by T2 with a phase delay of approximately 120°, then T1 with another 120° delay. These delays were largely consistent between legs and across animals, independent of speed (Fig. 3H). In fast animals, many legs were significantly coherent with each other (Fig. 3J). In slower animals, average cycle period was 0.7 ± 0.1s, and some legs often swung alone during walk cycles (Fig. 3E). Furthermore, not all six legs were required to swing during each cycle. Absent leg swings consequently impaired intrasegmental coordination; phase differences between contralateral leg pairs increased (binned mean for 5 slowest larvae = 41.0 ± 7.5°) compared to faster animals (binned mean for 5 fastest larvae = 15.1 ± 3.4°) (Fig. 3I). As such, fewer legs were significantly coherent to each other in slower animals (Fig. 3J), but the T3 to T1 sequence was largely preserved (Fig. 3H). Finally, analysis of all swing and stance phases against cycle period showed stance duration increased with cycle period, whilst swing duration was constant (Fig. 3K). Therefore, swing and stance duty cycles, defined as the proportion of time normalized to the cycle period, showed opposing exponential relationships with cycle period (Fig. 3L).

### Thoracic-abdominal coordination is crucial for stable, effective locomotion

To begin to understand the neural control of movement in *Tribolium* larvae, we severed the connective between thoracic and abdominal ganglia (Fig. 4A,B) to assess the effects of interrupting ascending and descending connections between brain and suboesophageal ganglion (SOG) and ventral nerve cord (VNC) on unrestrained locomotion. Severed animals travelled significantly less far (mean = 5.83 ± 0.58cm) on paper than controls (mean = 13.3 ± 2.0cm, p = 0.020) and sham treated animals (mean = 14.4 ± 2.0cm, p = .008; Fig. 4C,D). Moreover, they displayed a narrower range of body bending angles (range = 115.8 to 224.1°) than controls (range = 122.7 to 245.4°) and shams (range = 111.0 to 273.2°) (Fig. 4E). To see how these impairments affected performance during a locomotor challenge, we first performed preliminary experiments gauging control larvae performance on different inclinations: 180° (flat surface), 135° (uphill), 90° (vertical wall), and 45° (overhang) (Fig. S1). We found control animals walked progressively less far with greater deviations from 180° but were still able to cling onto and climb 45° overhangs. Interestingly, larval paths became straighter, and fewer animals approached arena edges with greater inclination (Fig. S1A-D,F).

**Figure 4:**
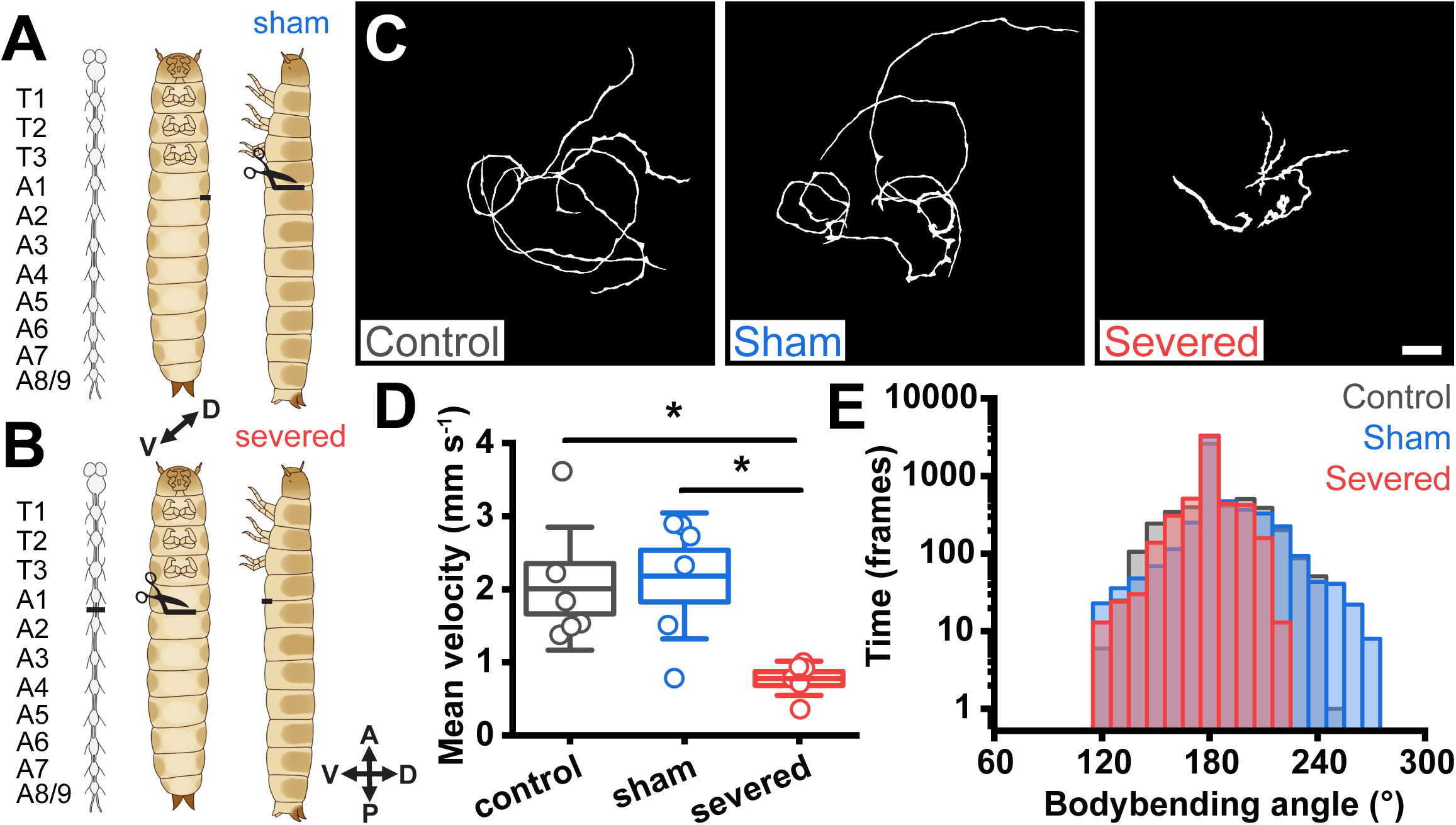
Severing thoracic-abdominal connectives impairs locomotion and restricts bending. (A,B) Surgical manipulations for (A) sham: lateral lesion between A1-A2 and (B) severed: ventral severing of the A1-A2 connectives. (C) Walk paths on paper across treatments for 1 minute. Scale bar = 2cm. (D) Mean instantaneous velocity across treatments. Boxes = mean ±SE, whiskers = ±SD; * p < .05, One-way ANOVA, n = 6. (E) Frequency plot of all body angles throughout 1 minute of locomotion at 15fps, binned every 10°. Note the exponential y axis (log10).

**Figure 5:**
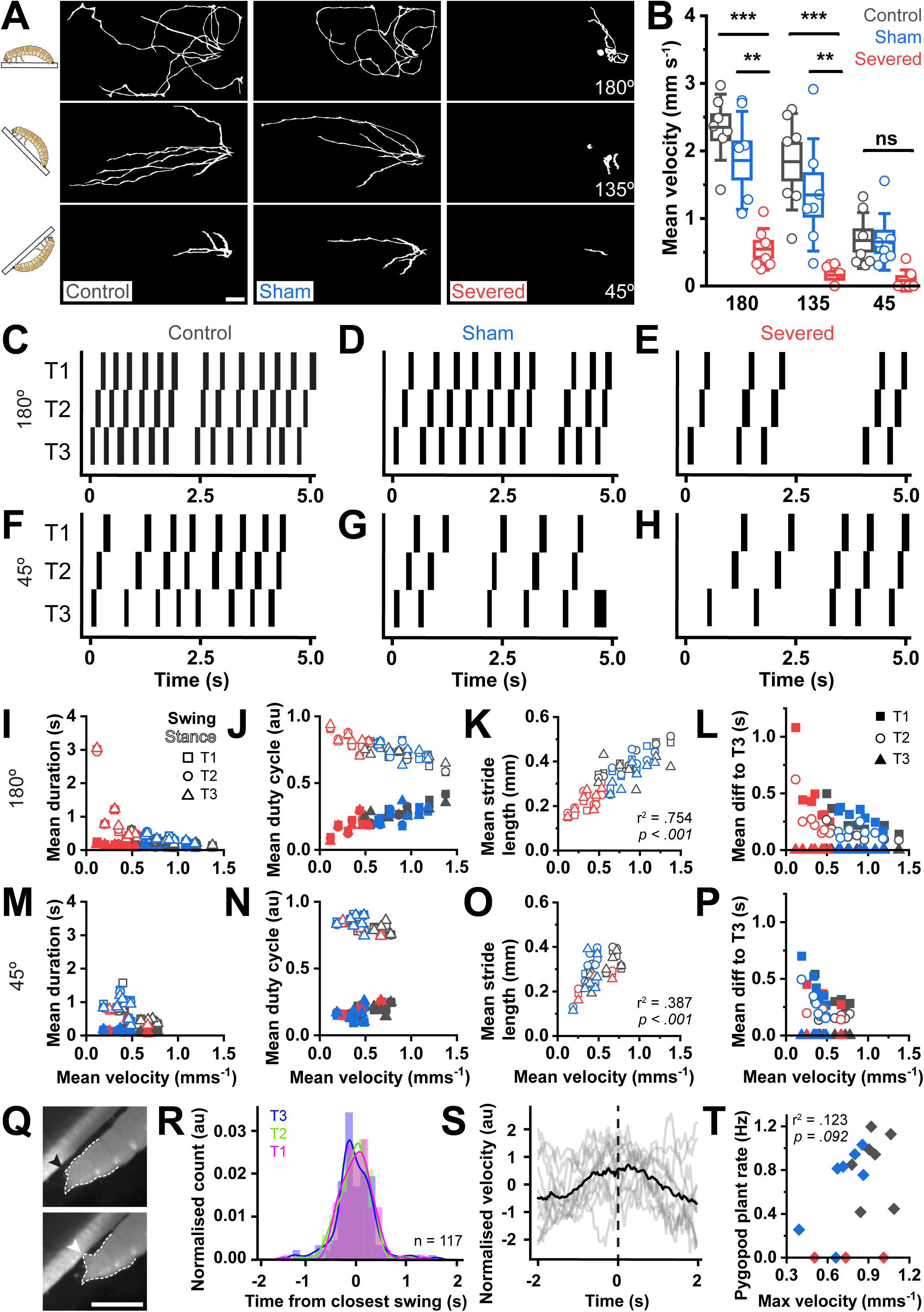
Abdominal inputs are not required for thoracic coordination and pygopods provide stability in locomotion. (A) Walk paths of control, sham, and severed larvae on top-down incline rig. Scale bar = 2cm. (B) Mean instantaneous velocity across treatments and inclinations. Boxes = mean ±SE, whiskers = ±SD; *** p < .001, ** p < .01, One-way ANOVA, n = 7. (C-H) Representative swing (black) and stance (white) phases of walk cycles following T3 to T1 sequence on up-close incline rig. Note the prolonged stance phases in severed at baseline 180°. (I-P) Scatter plots comparing locomotor parameters across surgical manipulations for 180° and 45°: (I,M) swing (filled shapes) and stance (empty shapes) duration, (J,N) swing and stance duty cycle, (K,O) stride length and (L,P) time difference to T3 (intersegmental difference). All values are averaged per segment (squares, circles, triangles); velocity is averaged per animal. (K,O) Stride length correlates significantly with velocity at 180° (r^2^ = .754, p < .001), and 45° (r^2^ = .387, p < .001), pooled across all animals. (Q) Frames showing pygopod relaxation parallel to substrate (black arrow) and planting into substrate (white arrow). Scale bar = 1mm. (R) Histogram of normalized pygopod planting event count relative to initiation of closest leg swing across all walk cycles, binned according to the Freedman–Diaconis rule (see Methods). Note the order of distribution peaks highlights predominant T3 to T1 gait even in unfiltered walk cycles. Total number of pygopod plant events analyzed = 117 pooled across 13 larvae. (S) Normalized mean instantaneous velocity (black trace) relative to pygopod planting events (dotted line); individual velocities across 13 larvae (control and sham combined) in grey. (T) Pygopod plant rate correlates insignificantly and weakly with maximum velocity (r^2^ = .123, p = .092), pooled across all animals. N = 7 per treatment.

Next, we used inclines as a motor challenge for severed animals to investigate how disconnecting the abdominal nervous system would affect clinging and climbing during challenged locomotion. Control, sham, and severed animals were tested on 180°, 135° and 45°. Severed animals exhibited prominent clinging and climbing impairments – 42.9% at 135° and 71.4% at 45° of larvae failed to cling on and fell (represented as 0mm s^-1^ points for velocity in Fig. 5B). Severed animals that could cling and climb seemed to have straightening paths and had slower speeds with increasing incline, as was observed in control and sham treated animals (Fig. 5A-H, Fig. S1G). As expected, severed speeds were significantly lower for baseline 180° (control-severed, p < .001; sham-severed, p = .002) and 135° (control-severed, p < .001; sham-severed, p = .007). Although some severed animals had curved postural phenotypes, this was unrelated to their speed of locomotion (curved individuals ranked 3^rd^, 4^th^ and 5^th^ out of 7 total severed larvae in speed). Nonsignificant differences at 45° may reflect an overall speed limitation at such extreme inclinations.

To assess leg kinematics and coordination differences across treatments and inclinations, we observed locomotor performance on 180° and 45° inclines on an up-close incline behavioral rig as opposed to the top-down view <u>(Movie 3)</u>. At 45°, 14.3% of severed animals fell, and 42.9% clung on but remained immobile. Consequently, these animals were excluded from locomotor and kinematic analysis. Notably, no control or sham animals fell at any inclination in either experiment. To effectively compare locomotion across conditions, walk cycles following the T3 to T1 gait sequence were analyzed. Severed animals were slower overall but followed locomotor parameter trends in control and sham for 180°. However, they were indistinguishable from controls and shams at 45° as overall locomotion slowed. Swing/stance durations and duty cycles followed the same trends with speed observed previously (Fig. 3K-L) independent of inclination; stance duration was modulated, whilst swing duration remained constant across speeds (Fig. 5I-J,M-N). Stride lengths increased linearly with speed at both 180° (r^2^ = .754, p < .001; Fig. 5K) and 45° (r^2^ = .387, p < .001; Fig. 5O), and intersegmental differences were proportional across treatments and inclinations (Fig. 5L,P).

At 45°, only control and sham larvae engaged their pygopods to assist climbing <u>(Movie</u> <u>3)</u>, which suggested climbing required descending inputs to the abdomen. Moreover, larvae consistently attempted to maintain abdominal contact with the substrate, suggesting sustained contraction of body wall muscles whilst climbing. Pygopod planting events were defined as visible pygopod extensions, and a change from a parallel alignment with the substrate to a perpendicular pointed orientation into the substrate (Fig. 5Q). Analysis of their relation to leg kinematics in all walk cycles showed that most pygopod planting events coincided temporally with the initiation of leg swing phases (Fig. 5R), and pygopod planting-triggered analysis showed the sequence from T3 (blue) to T1 (magenta) as seen by the peaks in the distributions (Fig. 5R). Moreover, pygopod engagement was preceded by an increase in velocity and followed by a decrease (Fig. 5S), consistent with the association between pygopod planting events and leg swings. Although most control and sham larvae engaged their pygopods on the overhang (mean rate = 0.76Hz, range = 0 to 1.20Hz), this did not significantly correlate with their maximum velocity (r^2^ = .123, p = .092) (Fig. 5T).

### Abdominal inputs are essential for tunnelling

As evidenced, abdominal inputs (ascending or descending) facilitated free locomotion on flat surfaces and during locomotor challenges. Although clinging and climbing behavior may assist *Tribolium* in navigating and infesting food storage systems, we wanted to probe the effects of severing thoracic-abdominal connectives on tunnelling into their food source. Larvae (control, sham and severed) were individually dropped onto a dish filled with approximately 10mm deep whole wheat flour and visible body length was measured 0s and 120s afterwards (Fig. 6A-F). Control and sham animals immediately tunneled into flour on contact, as observed previously during substrate preference tests, with 100% of control and 75% of sham larvae being fully submerged after 2 minutes (Fig. 6G). Strikingly, 0% of severed animals tunneled effectively into flour, and on average, 90.7% of their original body lengths was exposed after 2 minutes (Fig. 6H). Severed animals still visibly displaced flour with their legs, albeit without making any progress (Fig. 6F). Therefore, the abdominal segments are essential for larval submergence into flour.

**Figure 6:**
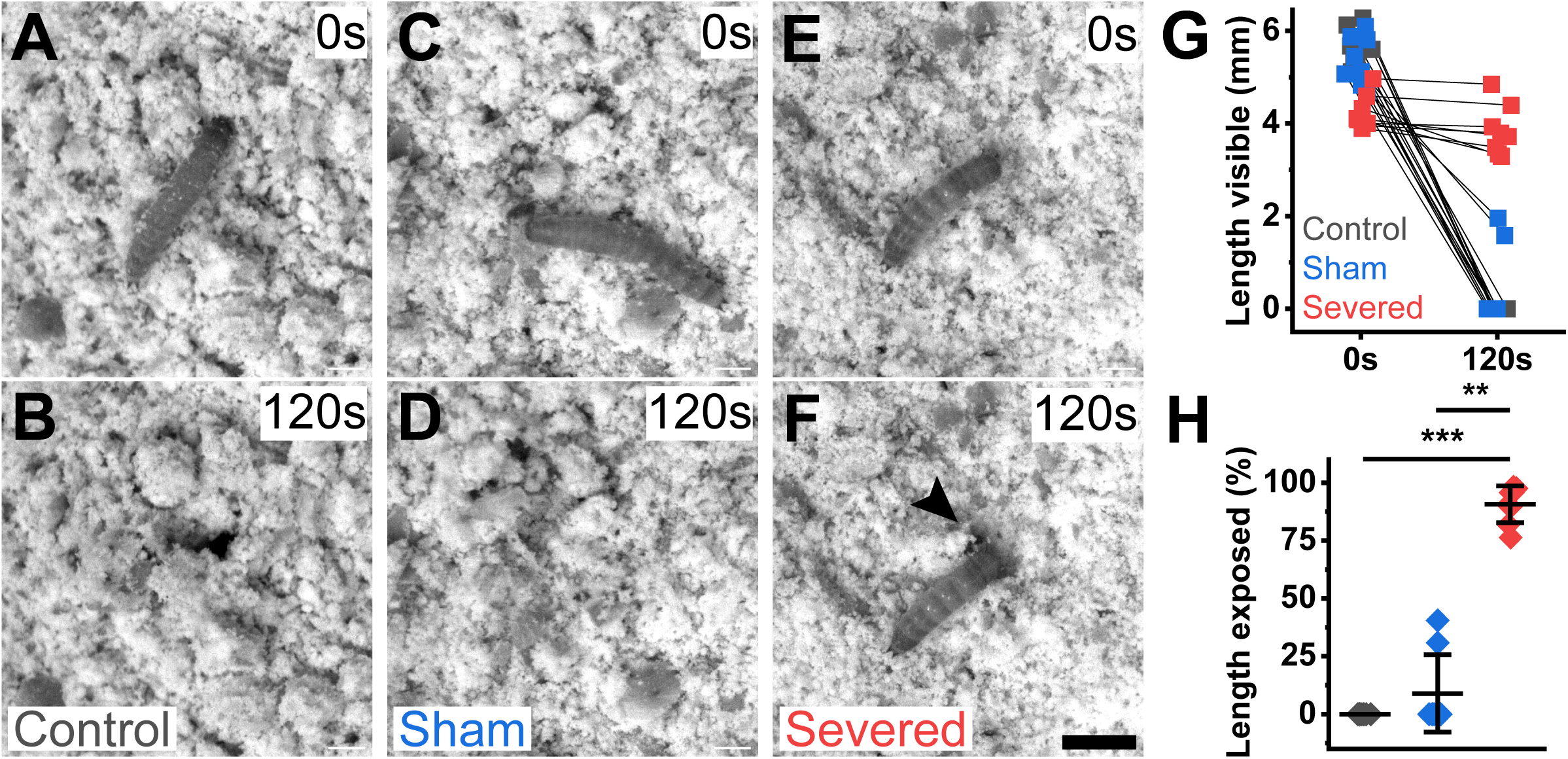
Abdominal inputs are crucial for larval tunnelling into natural food source. (A-F) Top-down frames at 0s and 120s after placement on flour for (A,B) control, (C,D) sham and (E,F) severed animals. Severed animals struggle to tunnel; black arrow in (F) indicates displacement of flour from thoracic leg movements despite no body submergence. Scale bar = 2mm. (G) Change in raw body length between 0s and 120s across treatments. (H) Percentage of original body length exposed after 120s. Whiskers = mean ±SE; *** p < .001, ** p < .01, Kruskal-Wallis test. N = 8 per treatment.

## Discussion

To study and understand the *Tribolium* larval abdomen’s roles in locomotion and its relationship to the legs, we first characterized the larva’s general locomotor performance across a range of conditions. We found larvae relied on their legs for unchallenging locomotion on preferred substrates, which mostly consisted of fibrous materials. This supports the hypothesis that their original habitat consisted of tree bark (Dawson, 1977; Good, 1936), and explains why wet, slippery surfaces present a challenge. While adults are capable of locomotion across flour (Pai, 2010), tunnelling is the primary motor program for navigating their food source during pre-pupal instars. This behavior reflects ethologically relevant photophobic preferences (Duehl et al., 2011) and their propensity to tunnel deeper when approaching pupation (King and Dawson, 1972). The locomotor patterns underlying their exploratory behavior consist of straight runs and continuous turning, with little to no periods of immobility. This differs from *Drosophila* larvae, which combine repeated straight runs with pauses followed by headsweeps, redirecting them to a new trajectory (Lahiri et al., 2011; Suster and Bate, 2002). *Tribolium* larvae also show no evidence of handedness in their exploratory movements, again in contrast with *Drosophila* adults (Buchanan et al., 2015) and larvae (Wosniack et al., 2022) but also other insects such as ants (Hunt et al., 2014) and cockroaches (Camhi and Johnson, 1999). This may reflect the requirements for *Tribolium* to navigate a 3-dimensional environment of constantly shifting substrate. We also found a larval preference for edge exploration, which may represent photophobicity, centrophobicity or a thigmotactic drive to maximize body contact with their environment to mimic how they are naturally enveloped in substrate. This behavior is also found in *Drosophila* adults, but not for centrophobic or thigmotactic reasons (Soibam et al., 2012).

The most common gait in insects is the alternating tripod gait, in which the first and third leg on one side of the body move together with the contralateral second leg (Wilson, 1966). *Tribolium* larvae, however, use a travelling wave gait, involving posterior-to-anterior sequences of swings propagating from T3 to T1 across all walking speeds. Their contralateral leg pairs move synchronously, and phase delays between segments are largely maintained across speeds. This bilaterally symmetric travelling wave gait is found in the larvae of other beetles, including leaf beetles (Gustafson and Chaboo, 2009; Zurek et al., 2015a). Since environmental conditions do not differ between larval and adult stages in these animals, the wave gait could therefore reflect developmental constraints on the circuit architecture driving locomotion in the Cucujiformia infraorder of beetles. However, the travelling wave gait pattern is found more widely. For example, in krill and crayfish, it generates the forces required for locomotion in an aquatic medium (Mulloney and Smarandache-Wellmann, 2012; Zhang et al., 2014), and in some millipedes it is used to generate the required forces for burrowing in soil (Manton, 1961). A similar bilaterally symmetrical gait has been observed in tardigrades, which is employed in a substrate-dependent manner.

Tardigrades more commonly use a tetrapod gait, but transition to a wave gait on softer and therefore unpredictable surfaces (Nirody et al., 2021). Similarly, the tobacco hornworm caterpillar, *Manduca sexta,* usually adopts a “progressive” posterior-to-anterior wave gait during horizontal crawling on stiff substrates, like *Tribolium* larvae, but transition to “non-progressive” gaits on soft substates (Metallo et al., 2020). Another example is the dung beetle *Pachysoma endroedyi* ‘galloping’ gait, in which the anterior two leg pairs move in a travelling wave, with the hind leg pair dragging behind them and contributing little to forwards motion (Smolka et al., 2013). In these desert dwelling species, in-phase contralateral coordination may be energetically favorable for traversing constantly shifting terrains like sand, a constraint they might share with aquatic species. These considerations are equally relevant for *Tribolium,* as they consume and inhabit a constantly deformable substrate. This gait may become particularly beneficial during *Tribolium* pre-pupal stages, when larvae tunnel deeper for warmth and lower population densities to prepare for pupation (King and Dawson, 1972).

Gait adjustments tune movement to the requirements of different terrains and speeds of locomotion (Bässler and Büschges, 1998; Bender et al., 2011; Graham, 1972; Ramdya et al., 2017; Schilling et al., 2013; Smolka et al., 2013). Many animals display speed-dependent gait transitions. For example, adult *Drosophila* gaits have been shown to transition from a pentapodal wave (one leg is in swing phase at a time) to a tetrapodal (two legs, but not bilateral pairs) and finally tripodal wave (three legs) by modulating stance phases as the animal progresses from slow to fast speeds (DeAngelis et al., 2019). We therefore also characterized the kinematic changes accompanying different speeds of locomotion in *Tribolium* larvae. They similarly have increasing numbers of limbs in swing phase at a time during increasingly fast locomotion, but do not appear to adopt a gait other than their bilaterally symmetrical metachronal wave. In addition, *Tribolium* larvae increase walking speeds by reducing stance duration. This is a widely observed locomotor strategy used across phyla, from humans (Grillner et al., 1979), to cats (Frigon et al., 2014), to *Drosophila* (DeAngelis et al., 2019) and tardigrades (Nirody et al., 2021). Moreover, larval stride length increases linearly with speed, also adhering to a highly conserved locomotor strategy across invertebrates and vertebrates (Granatosky and McElroy, 2022; Hooper, 2012; Strang and Steudel, 1990). A similar in-depth analysis of adult *Tribolium* gait kinematics in the future will allow further comparison.

The *Tribolium* VNC is unfused, with a separate ganglion in each body segment, chained together by longitudinal connectives, and each thoracic ganglion innervating a leg pair (Farnworth et al., 2022a; Hunnekuhl et al., 2020). This anatomical arrangement is found across arthropods (Richter et al., 2010) and tardigrades (Mayer et al., 2013), and suggests gait variation across and within species might be derived from one single modifiable control strategy (Nirody, 2021). This raises the question whether the neural control of its behavior shares similarities with that of the more well-studied and soft-bodied *Drosophila* larva, which is legless and moves by means of abdominal peristaltic waves (Lahiri et al., 2011), and has a segmentally fused central nervous system (Truman and Bate, 1988).

Qualitatively observed pygopod planting during unrestricted motor programs (backtrack and redirect, see Fig. 2F) first suggested a prominent role for the abdomen in assisting natural movement. We present several lines of evidence that pygopods act to support locomotion in challenging conditions. First, while fast animals crawling on level fibrous surfaces do not necessitate pygopod engagement, they do while climbing 45° overhangs. Furthermore, pygopod planting events are largely temporally locked to leg swing phases, during which the animal needs to maintain static stability, and these events are preceded by a temporary increase in crawling speed. This shows a correlation between situations that compromise the animal’s stability and likelihood of pygopod engagement. Furthermore, the weak, non-significant correlation between pygopod planting rate and maximum velocity further implicates its role in stability over a mechanism to boost/propel overall speed. Second, severing the A1-A2 connective, thus removing descending and ascending inputs between the brain, thorax and abdominal segments, resulted in lower speeds during walking and climbing. Some severed animals dragged their abdomen during walking <u>(Movie 3)</u>, which may contribute to decreasing velocity as the lack of abdominal sensory inputs may impair its ability to support itself during locomotion. However, impairments on inclines varied, ranging from some severed animals falling off mild (135°) inclines to a select few matching control and sham performance on 45° overhangs. The latter may reflect the overhang limiting the feasible speeds larvae can produce in this motor challenge. Like the variability in inter-animal velocities in normal larvae, the severity of severing is likely equally dependent on the individual. Nevertheless, severed animals never engaged their pygopods, and most tended to fall off 45° overhangs. These animals’ inability to engage their posterior segments therefore particularly impaired their locomotor capabilities under challenging conditions. Furthermore, this phenotype is unlikely due to a failure to coordinate leg movements, since severed animals fall into the same speed-dependent distributions of kinematic patterns including swing and stance duration, duty cycle, stride length and intrasegmental phase difference as control and sham larvae. This suggests that abdominal regions are not required for fundamental leg coordination but provide a crucial support role in larvae.

Other species that have abdominal anchoring points with potential stabilizing roles include *Labidomera clivicollis* larvae, which use pygopods during body extension whilst climbing tree branches (Gustafson and Chaboo, 2009); and blackfly larvae, which are adapted to anchor into riverbeds to avoid getting swept away by strong currents (Jamnback and Frempong-Boadu, 1966). The *Tribolium* larval anatomical adaptations we describe that confer stability and allow the animal to stay upright may similarly help evade predation, for example by the parasitoid wasp *Holepyris sylvanidis* (Amante et al., 2018), either by enhancing locomotion efficiency in shifting terrain or avoiding falling off tree branches in their original habitat. The use of posterior terminal organs for substrate engagement during locomotion is also found in other soft-bodied animals including *Drosophila* larvae (Booth et al., 2024), *Gastrophysa viridula*, which deploy their pygopods phase-locked to leg movements during the locomotor cycle (Zurek et al., 2015b); and *Manduca sexta*, whose locomotor cycle starts with the terminal prolegs lifting off the substrate (van Griethuijsen and Trimmer, 2014). Our results elucidate a functional role for the pygopods, which were previously only phenotypically mentioned in the *iBeetle* RNA interference (RNAi) screens (Hakeemi et al., 2022; Schmitt-Engel et al., 2015).

The contribution of abdominal segments is likely to extend beyond offering physical support during locomotion. During motor challenges like overhangs, larvae may send ascending information from abdominal sensory neurons (e.g., regarding reduced body contact with substrate) to sensorimotor integration centers for sensory feedback. These may then integrate and send back descending information regarding internal representations (e.g., current geotactic location, motor state) to make a higher order decision to plant the pygopods to increase stability. Interruption of communications to and from the abdomen and thus lack of stability in severed animals may explain those that fall off or fail to climb. However, the severed animals that could cling and climb followed the same leg kinematics as controls and shams, therefore lack of abdominal input appears to not disrupt thoracic movements.

Furthermore, our data suggest the posterior segments have prominent functions in tunnelling alongside stability. Tunnelling is a key motor program for navigation, feeding, development, and reproduction in their natural habitat, and the inability to tunnel in severed larvae suggests a crucial role for abdominal contractions, pygopod recruitment, and/or ascending sensory feedback in coordinating effective tunnelling. The abdomen may be recruited to generate pushing forces to submerge larvae in flour, similar to millipedes (Manton, 1961). Equally, it may aid exploratory movements such as the backtrack and redirect behavior (Fig. 2F) to search for new food sources. Investigating the structure of both ascending and descending inputs passing through the larval *Tribolium* A1-A2 connective is therefore a promising avenue to explore the neural basis of its behavior and could be a possible pinch point to disrupt the animal’s ability to infest grain stores.

Finally, *Tribolium* is an attractive emerging model system in behavioral neuroscience, combining a robust experimental toolkit with an extensive body of work describing its development and physiology. As the first coleopteran species to have a fully sequenced genome (∼16,500 genes), 47% of which are orthologous to insects and vertebrates, *Tribolium* are already highly favored in the field of developmental genetics (Richards et al., 2008). Their genetic toolkit includes the GAL4-UAS system (Schinko et al., 2010), CRISPR/Cas9 genome editing (Adrianos et al., 2018; Gilles et al., 2015), balancer chromosomes, and stable injectable RNAi gene silencing (Tomoyasu et al., 2008). Forward genetic screens uncovered previously unknown genetic regulators of leg and body wall muscle development (Schultheis et al., 2019), and recent studies have described the structure and function of the *Tribolium* cryptonephridial system, which is essential for the animal’s exceptional tolerance to desiccation (Naseem et al., 2023). The structures and locations of brain regions in larval and adult *Tribolium* showed broad similarities to other insect model organisms (Farnworth et al., 2022b; Garcia-Perez et al., 2021; He et al., 2019). In addition, comparison of *Tribolium* and *Drosophila* larval neuromuscular systems revealed resemblances in somatic muscle composition (Schultheis et al., 2019). Its similarities to both ancestral and highly derived insects therefore make it an attractive comparative model.

Overall, this study has emphasized how the kinematic analysis of movement and underlying neural control provides insights into natural behaviors that govern *Tribolium*’s success as a pest. We identified the importance of abdominal segments and pygopods to both locomotor challenges and tunnelling, suggesting that studying the structure and function of its connectives has important potential implications for behavioral neuroscience and food security. How far our findings extend to earlier instars and adults remains a promising question for future study.

## Acknowledgements

We thank Barry Denholm for providing *Tribolium* stocks, Adam Taylor for *Groundswell*, Balwant Singh for building the up-close incline rig, Chris Sutherland for statistical advice and Pulver Lab and Zwart Lab members for constant support.

## Author contributions

Conceptualization: B.X.Y., M.F.Z. and S.R.P.; Methodology: B.X.Y.; Investigation: B.X.Y.; Formal Analysis: B.X.Y. and M.F.Z.; Visualization: B.X.Y.; Writing: B.X.Y., M.F.Z. and S.R.P; Funding Acquisition: B.X.Y.; Supervision: M.F.Z. and S.R.P.

## Data and resource availability

Raw experimental movies are available at the University of St Andrews Research Portal: doi:10.17630/a97be752-0c97-42a2-ada4-745c5acff342. Analyzed files are available at Zenodo: doi:10.5281/zenodo.11520087. Analysis code for gait and incline results are available upon request.

---

**Figure S1:**
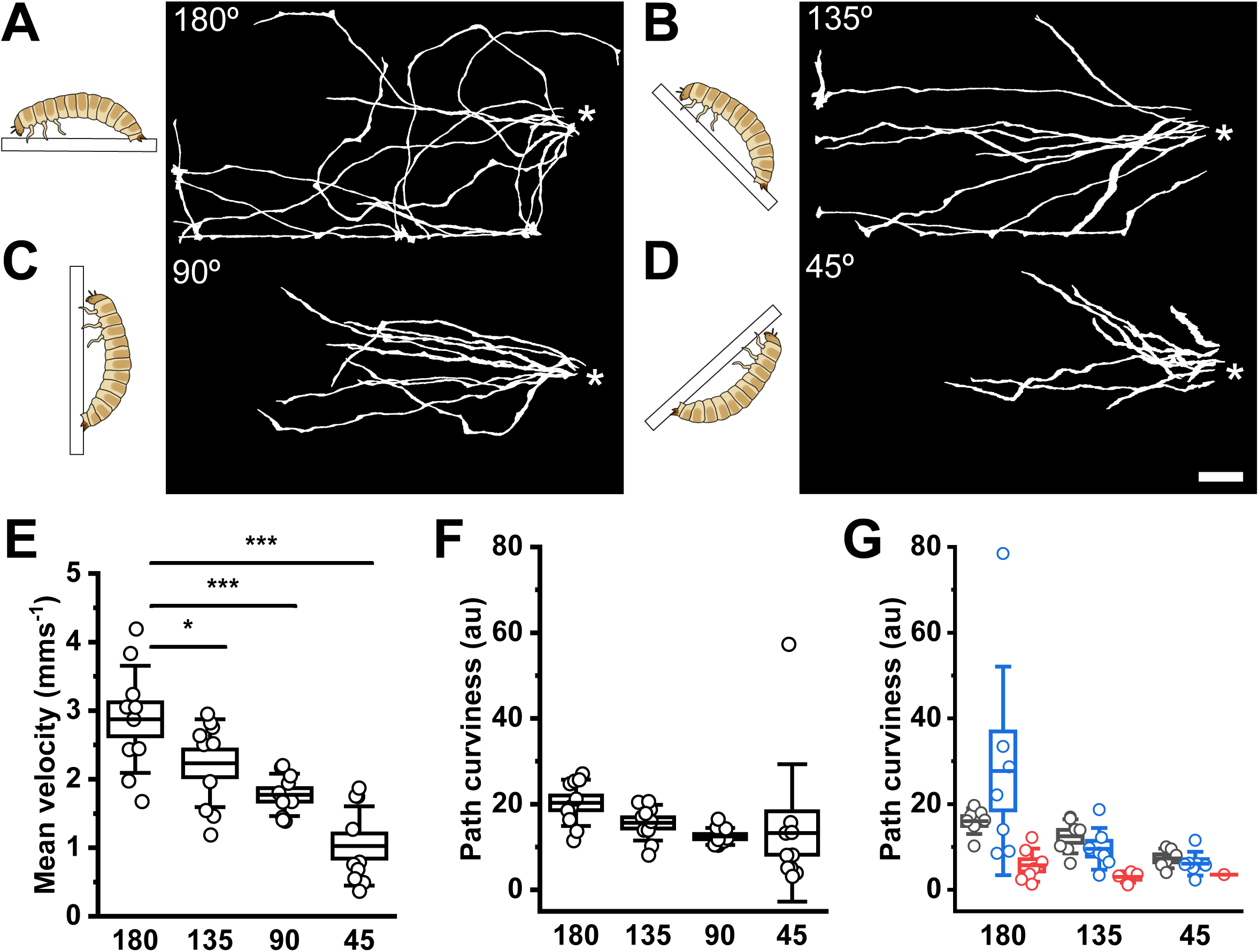
Increasing incline becomes progressive locomotor challenge for *Tribolium* larvae. (A-D) Left: diagrams of larval orientations relative to angled platforms (white rectangles). Right: walk paths of larvae on respective platforms. Note, 45° is an overhang. Asterisks indicate starting position. (E) Mean instantaneous velocity across inclinations. Boxes = mean ±SE, whiskers = SD; *** p<.001, * p<.05, One-way ANOVA, n = 10. (F) Path curviness across inclinations calculated via (total distance / displacement) * max distance from start for preliminary control trials in (A-E), n = 10. (G) Path curviness for experimental trials in Fig. 5A,B across inclinations and surgical manipulations, n = 7.

